# Perivascular space mediated the interaction between sleep, and brain functional connectivity in the healthy aging population

**DOI:** 10.1101/2025.05.19.654921

**Authors:** Nien-Chu Shih, Joey Contreras, Wendy J. Mack, Jeiran Choupan

**Affiliations:** Laboratory of Neuro Imaging, USC Mark and Stevens Neuroimaging and Informatics Institute, Keck School of Medicine, University of Southern California, Los Angeles, CA, USA; Alzheimer’s Disease Cooperative Study (ADCS), University of California San Diego, La Jolla, CA, USA; Department of Population and Public Health Sciences, Keck School of Medicine, University of Southern California, Los Angeles, CA, USA; NeuroScope Inc., New York, USA

**Keywords:** Perivascular space, Functional connectivity, Magnetic resonance imaging, Sleep, Cognition

## Abstract

Perivascular space (PVS) surrounds the perforating arteries or draining veins of the cerebral cortex as part of the brain clearance system. Previous studies showed that sleep affects both brain clearance function and brain functional connectivity (FC). However, the impact of PVS characteristics on brain FC remains unclear. This study investigated these associations and their link to cognition. We utilized cross-sectional structural MRI and resting state-fMRI data from 512 health aging population in the HCP-Aging dataset, together with Pittsburgh Sleep Quality Index questionnaire and NIH cognitive tests. Our results showed that basal ganglia (BG)-PVS volume fraction (VF) was positively correlated with FC in the right anterior medial temporal gyrus (aMTG) and right temporal regions, while centrum-semiovale (CSO)-PVS VF was positively correlated with FC in the left hippocampus and right frontal regions. Increased CSO-PVS VF in early middle-aged adults showed higher hippocampal FC and better cognitive performance. Interestingly, individuals with longer time spent in bed had larger BG-PVS VF linked to higher FC in the right aMTG. Additionally, older adults with better sleep quality had larger BG-PVS VF linked to higher FC in the right aMTG. These findings suggest that PVS morphology may reflect changes in neural connections involved in memory-related regions.

## 1 Introduction

The brain clearance system is a highly specialized fluid transport systems that facilitate the exchange of interstitial fluid and regulates cerebrospinal fluid (CSF) dynamics within the brain^1^. Dysfunction of the brain clearance system has been associated with decline in cognitive function and degeneration of nerve cells^2^. Previous studies have shown that the brain clearance system may play a role in the development and progression of neurodegenerative disorders such as Alzheimer’s disease (AD) and Parkinson’s disease^1^. Perivascular spaces (PVS), an important segment of the brain clearance system, are CSF-filled spaces surrounding small blood vessels in the brain and play a crucial role in brain clearance of amyloid-β and other metabolic waste ^3,4^. Previous research suggests that waste clearance occurs most readily during deep sleep, as evidenced by the movement of molecular tracers from the cisterna magna CSF, along periarterial spaces, and through brain parenchyma^5^. Our team have made attempts to investigate PVS in cognitive decline and the link between PVS and sleep. Among all the results, we notably observed reduction in PVS volume fraction (VF), measured by structural MRI, in the anterior superior medial temporal gyrus (asMTG) of individuals with mild cognitive impairment when compared to cognitively normal aging controls^6^. In a separate study, to our surprise, we found that healthy older adults who had better sleep quality had larger PVS VF^7^. To better understand the brain clearance mechanism, this study aims to investigate how poor sleep and structural changes in PVS influence brain functional neuronal activity.

In aging and neurodegenerative diseases like AD, disruptions in sleep are often accompanied by altered functional connectivity (FC) within the Default Mode Network (DMN)^8^, assessed by analyzing Functional Magnetic Resonance Imaging (fMRI) data. The DMN involves in self-referential thinking, memory consolidation, and internal cognition, usually active at rest and inactive at work^9^. fMRI utilizes the Blood Oxygen Level Dependent (BOLD) signal as an indirect measure of neuronal activity^10,11^. The BOLD signal detects changes in the ratio of oxygenated to deoxygenated hemoglobin in the blood^10^. When neuronal activity leads to increased metabolic demand, it promotes a localized increase in cerebral blood flow (CBF) that overcompensates for oxygen consumption, resulting in a relative decrease in deoxygenated hemoglobin and a corresponding BOLD signal change^11^. In addition, early amyloid-β deposition may affect DMN connections of cognitively healthy individuals^12,13^. Furthermore, long-term sleep deprivation has been shown to decrease DMN activity and connectivity, with poor sleep quality similarly associated with weaker DMN function^14,15^. Notably, previous studies have indicated that sleep deprivation upregulates connectivity in the DMN and impairs the frontal–parietal functional network^16,17^. Moreover, sleep disorders such as insomnia and daytime sleepiness, result in reduction and disturbance of DMN functional connectivity^18,19^. These findings highlight that sleep behavior may influence cognitive performance by altering brain connectivity patterns, and that sleep quality and patterns play an integral role in modulating brain activity.

Despite findings on how sleep affects both brain clearance systems and DMN function, the link between sleep behavior, PVS and FC in healthy individuals remains understudied. Therefore, in this study, we conducted an examination on resting state FC using the Human Connectome Project Aging (HCP-Aging)^20^ dataset to gain deeper insights into the link between PVS, sleep, and FC in healthy individuals.

## 2 Material and Methods

### 2.1 Participants

Structural MRI and resting state functional MRI (fMRI) data from 512 cognitively healthy individuals aged 36-100 years from the HCP-Aging Lifespan Release 2.0^20^ were examined. Participants had no diagnosed neurological or major psychiatric disorders at recruitment. In this study, for each participant we used their information on age, sex, years of education, Pittsburgh Sleep Quality Index (PSQI) questionnaire, cognitive performance measured by the Montreal Cognitive Assessment (MoCA), NIH toolbox working memory tests, and NIH toolbox cognition battery (total cognition composite score, total fluid cognition composite score, and crystallized cognition composite score).

While the dataset excluded individuals with diagnosed conditions, it did not exclude based on sleep behavior. MoCA scores between 20 and 30 were classified as healthy aging despite scores below 25 typically reflecting mild cognitive impairment. All exclusion criteria for the current analyses on HCP-Aging data are following our previous study^21^. We further excluded one participant who had missing fMRI data. Therefore, 512 participants were included in the analyses. We followed the cutoff age range buckets based on previous reserach^22–25^,we categorized the participants by age as early middle-aged (36-55 years old, N=252), middle-aged adults (56–65 years old, N=111), or older adults (above 65 years old, N=149). The demographic information of the participants is shown in Table 1.

**Table 1.** Demographic information of the study cohort.

| Characteristic | Overall<br>N = 512 | Early Middle-aged<br>(age 36-54), N =<br>252 | Middle-aged<br>(age 55-65), N =<br>111 | Older<br>(age >65), N =<br>150 | p-value<sup>3</sup> |
| --- | --- | --- | --- | --- | --- |
| **General** |  |  |  |  |  |
| **Age (yrs)<sup>1</sup>** | 55 (45, 67) | 45 (41, 49) | 59 (57, 62) | 73 (69, 79) | <0.001** |
| **Sex<sup>1</sup>: Male** | 215 (42%) | 97 (38%) | 50 (45%) | 68 (45%) | 0.3 |
| **Female** | 297(58%) | 155(62%) | 61(55%) | 82(55%) |  |
| **BMI (Kg/m<sup>2</sup>)<sup>1</sup>** | 26.4 (23.5, 30.2) | 26.6 (23.7, 30.8) | 25.5 (23.4, 28.5) | 26.5 (23.3, 30.2) | 0.4 |
| **Years of Education<sup>1</sup>** | 18 (16, 19) | 18 (16, 19) | 18 (16, 19) | 18 (16, 19) | 0.5 |
| **BG volume (mm3)<sup>1</sup>** | 38038 (36005, 40964) | 39694 (37391, 42233) | 37300 (35776, 39868) | 36296 (33962, 38666) | <0.001** |
| **CSO volume (mm3)<sup>1</sup>** | 245086 (223586, 266850) | 252153 (230731, 275540) | 244560 (224885, 265759) | 235571 (210592, 256462) | <0.001** |
| **BG PVS volume (mm3)<sup>1</sup>** | 768 (557, 1053) | 623 (479, 903) | 796 (671, 1039) | 1,032 (757, 1237) | <0.001** |
| **CSO PVS volume (mm3)<sup>1</sup>** | 5668 (4360, 7960) | 5037 (3803, 6962) | 6046 (4630, 8401) | 6281 (5149, 8807) | <0.001** |
| **BG PVS volume fraction (VF)<sup>1</sup>** | 0.0005 (0.0004, 0.0007) | 0.0004 (0.0003, 0.0006) | 0.0005 (0.0004, 0.0007) | 0.0007 (0.0005, 0.0009) | <0.001** |
| **CSO PVS volume fraction (VF)<sup>1</sup>** | 0.0039 (0.0030, 0.0053) | 0.0035 (0.0027, 0.0049) | 0.0040 (0.0032, 0.0058) | 0.0042 (0.0034, 0.0056) | <0.001** |
| **Cognition tests** |  |  |  |  |  |
| **MoCA score<sup>1</sup>** | 27 (25, 29) | 27 (25, 29) | 27 (25, 28) | 26 (25, 28) | 0.012* |
| **NIH Working memory score<sup>1</sup> (age-adjusted)** | 104 (93, 114) | 103 (91, 113) | 106 (92, 113) | 107 (96, 115) | 0.020* |
| **NIH total cognition composite<sup>1</sup> (age-adjusted)** | 110 (99, 120) | 109 (98, 121) | 108 (98, 119) | 111 (103, 118) | 0.7 |
| **NIH fluid cognition composite<sup>1</sup> (age-adjusted)** | 106 (95, 117) | 106 (93, 117) | 105 (94, 116) | 107 (98, 115) | 0.8 |
| **NIH crystallized cognition composite<sup>1</sup> (age-adjusted)** | 111 (100, 121) | 110 (99, 123) | 111 (102, 119) | 111 (105, 121) | 0.5 |
| **PSQI measurements** |  |  |  |  |  |
| **Sleep efficiency score<sup>2</sup>** | 0.44 ± 0.77 | 0.42 ± 0.71 | 0.46 ± 0.83 | 0.47 ± 0.80 | 0.2 |
| **Sleep quality score<sup>2</sup>** | 0.79 ± 0.67 | 0.84 ± 0.69 | 0.78 ± 0.71 | 0.70 ± 0.61 | 0.4 |
| **Sleep latency score<sup>2</sup>** | 0.82 ± 0.84 | 0.79 ± 0.85 | 0.85 ± 0.85 | 0.86 ± 0.83 | 0.7 |
| **Sleep medication score<sup>2</sup>** | 0.51± 0.99 | 0.34 ± 0.82 | 0.62 ± 1.05 | 0.71 ± 1.15 | 0.004* |
| **Sleep duration (hrs)<sup>1</sup>** | 7.0(1.5) | 7.0(1.5) | 7.0(1.5) | 7.0(1.44) | 0.045* |
| **Time in bed (hrs) <sup>1</sup>** | 8.0(1.5) | 7.8(1.5) | 8.0(1.5) | 8.0(1.8) | 0.10 |
| **PSQI total score <sup>2</sup>** | 4.63 ± 2.70 | 4.45 ± 2.68 | 4.80 ± 2.69 | 4.81 ± 2.72 | 0.4 |

<sup>1</sup> Median (IQR); n (%)
<sup>2</sup> Mean value ± standard deviation (SD)
<sup>3</sup> Kruskal-Wallis rank sum test; Pearson's Chi-squared test
\*Significant p-value < 0.05
\*\*Significant p-value < 0.001

### 2.2 MRI Acquisition and pre-processing

All participants were scanned using a customized Siemens 3T Prisma scanner housed at Washington University in St. Louis, using a standard 32-channel Siemens receive head coil. The T1-weighted image was acquired with repetition time (TR)/inversion time (TI) = 2500/1000 ms, time to echo (TE) = 1.8/3.6/5.4/7.2 ms, field of view (FOV) = 256 × 240 × 166 mm and the T2-weighted image with TR= 3200 ms, TE= 564 ms, FOV= 256 × 240 × 166 mm. Structural T1-weighted MPRAGE and T2-weighted SPACE images were preprocessed in LONI pipeline using the HCP minimal processing pipeline version 4.0.1 and FreeSurfer version 6. The preprocessing included gradient nonlinearity corrections, registering structural images together, aligning to native space anterior commissure-posterior commissure, and registering to MNI space using FSL’s FNIRT. Native space images were used to generate individual regional subcortical PVS features for white and pial surfaces with FreeSurfer. Detailed preprocessing methods are described in a prior publication^26^.

Resting state fMRI (rs-fMRI) scans were acquired with a 2D multiband (MB) gradient-recalled echo (GRE) echo-planar imaging (EPI) sequence (MB8, TR/TE=800/37 ms, flip angle=52°) and 2.0 mm isotropic voxels covering the whole brain (72 oblique-axial slices). Functional scans were acquired in pairs of two runs, with opposite phase encoding polarity so that the fMRI data in aggregate is not biased toward a particular phase encoding polarity. The HCP-A fMRI data used in this study were anterior-to-posterior (AP) and posterior-to-anterior (PA) phase encoding directions and were averaged through the default processing pipeline via CONN toolbox^27^ (release 20.b).

Participants acquired in total 26 min of resting state scans of four runs of 6.5 min each. During rs-fMRI scanning, participants view a small white fixation crosshair on a black background. Participants were instructed to stay still, stay awake, and blink normally while looking at the fixation crosshair.

### 2.3 PVS segmentation

Enlarged perivascular spaces (PVS) are more visible on basal ganglia (BG) and centrum semiovale (CSO) of MRI, therefore we chose these two areas as our PVS regions of interest (ROIs). The BG and CSO were segmented by FreeSurfer using an atlas-based approach, and brain volume and white matter masks were derived from the Desikan-Killiany atlas. PVS segmentation was performed as explained in Sepehrband et al^28^. In brief, T1w and T2w images were adaptatively filtered to remove the high-frequency noise and then co-registered and combined to obtain enhanced PVS contrast (EPC). EPC is shown to provide superior visibility of PVS compared with T1w or T2w alone (Supplementary Figure 1)^28^. PVSs were segmented from EPC images by applying Frangi filter using Quantitative Imaging Toolkit (QIT)^29^ and a vesselness threshold of 1.5 ^28^, which was optimized for the HCP data. To verify the accuracy of PVS segmentation, PVS segmentation quality control was performed by four trained analysts. Detailed quality control methods and evaluation criteria are described in a prior publication^28^.

### 2.4 Functional MRI preprocessing, denoising, analysis

All functional imaging data were processed with the CONN toolbox^27^. Functional and anatomical data were preprocessed via SPM^30^ (release 12.12.6) using a preprocessing pipeline^31^ including realignment with correction of susceptibility distortion interactions, slice timing correction, direct MNI-space normalization, and smoothing. The functional data were smoothed using spatial convolution with a Gaussian kernel of 6 mm full width half maximum (FWHM). Subsequently, functional data were denoised using a standard denoising pipeline^32^, which applies linear regression and bandpass frequency filtering to remove the effects of unwanted motion and other artifacts from the BOLD signal.

We conducted seed-to-voxel analysis to examine the association between PVS VF and the whole-brain FC of brain regions involved in DMN, bilateral anterior medial temporal gyrus (aMTG), bilateral parahippocampal gyrus and bilateral hippocampus. For the selection of seeds of functional connectivity, beside default mode network regions (medial prefrontal cortex, bilateral paracentral lobule, posterior cingulate cortex), we also examined hippocampus and parahippocampal gyrus, which are related to memory consolidation during the sleep^33^. In our prior research, we observed a notable reduction in PVS VF in the asMTG of individuals with MCI compared to cognitively normal aging cohorts^34^, therefore, we selected aMTG as one of regions of interest (ROI) regions. In this study, FC was defined as the correlation coefficients between the time series of specific seed regions and voxel clusters throughout the brain, reflecting the synchronization of neural activity across regions.

ROI FC maps were regressed on PVS VF in a general linear model, controlling for age and sex. Voxel-level threshold p < 0.001 and cluster-size p-FDR <0.05 were used, which were adjusted by FDR for multiple comparison correction.

### 2.5 Sleep parameters

The Pittsburgh Sleep Quality Index (PSQI) is a 19-item self-rated questionnaire used to assess subjective sleep quality over the past month^35^. Each item is scored from 0 to 3, with higher scores indicating poorer sleep. Sleep efficiency is calculated as the total sleep time divided by the time spent in bed, resulting in a score of 0 for a proportion greater than 85%, 1 for 75-84%, 2 for 65-74%, and 3 for less than 65%. Sleep quality is subjectively rated as very good (0), fairly good (1), fairly poor (2), or very poor (3). Sleep duration was calculated by subtracting the time participants fell asleep from the time they woke up. The duration of sleep is scored as 0 for more than 7 hours, 1 for 6-7 hours, 2 for 5-6 hours, and 3 for less than 5 hours. Sleep latency, or the time taken to fall asleep, is scored as 0 for less than 15 minutes, 1 for 16-30 minutes, 2 for 31-60 minutes, and 3 for more than 60 minutes. Sleep medication use is evaluated based on frequency, with scores of 0 for no usage, 1 for usage less than once a week, 2 for once or twice weekly, and 3 for three or more times per week. Daytime dysfunction, which assesses the ability to stay awake and maintain enthusiasm, is scored as 0 for no problems, 1 for very slight problems, 2 for somewhat of a problem, and 3 for a very big problem. Time in bed was calculated by subtracting the participants’ usual bedtime from their usual wake-up time. The PSQI total score, ranging from 0 to 21, is the sum of the above components, with higher scores indicating worse overall sleep quality.

### 2.6 Statistics

We used the fraction of the ROI volume that is occupied by the PVS (referred to as volume fraction) as our measure to calculate the PVS burden in the BG and the CSO. Owing to the non-normal distribution of the PVS in the BG, the log-transformed value of the VF of the PVS was used. The one-way analysis of variance (ANOVA) tests was used to compare sex, BMI, years of education, or cognitive performances between different age groups, and the results were shown in Table 1. FC values were extracted from the significant clusters obtained on CONN toolbox results.

We aimed to test whether (1) FCs were associated with PVS VF, (2) FC patterns that were significantly associated with PVS VF in the main analyzes were also correlated with global cognition and memory performance, and (3) the impact of sleep on PVS and these associated FC. Therefore, correlation analyses were performed between FC values and (1) PVS VF, (2) cognitive scores, and (3) sleep measurements in the whole sample and separately in early middle-aged, middle-aged, and older participants. Pearson Correlation was used for continuous variables, and spearman correlation was used for ordinal variables. Furthermore, to investigate if sleep impact the relationships between PVS VF and FC, we applied interaction testing on regression model.

To further explore the indirect effects of PVS on cognitive performance mediated through FC or the indirect effects of sleep on FC mediated through PVS, we utilized the mediation package in R to construct mediation model. A 95% confidence interval was reported to determine the significance of the indirect effects, with p < 0.05 indicating statistical significance. All regression and mediation analyses controlled for age, sex and years of education, and considered statistically significant at p-values < 0.05. FDR p-value was also performed for multiple comparison correction. In this study, multiple hypothesis tests were performed across PVS, FC, and sleep. The Benjamini-Hochberg procedure was applied at an alpha threshold of 0.05 to identify significant results while controlling for the proportion of false positives. All statistical analyses were conducted using R software (version 4.3.3).

## 3 Results

### 3.1 Clinical and Imaging Characteristics of Study Participants

Participant demographic and cognitive data are summarized in Table 1. In total, 252 early middle-aged participants (97 men and 155 women; age range 36–54 years), 111 middle-aged participants (50 men and 61 women; age range 55–65 years), and 150 older participants (68 men and 82 women; age range above 65 years) were included in this study. There were no significant differences in sex, BMI, years of education, or general cognitive performances (NIH total cognition composite score, fluid cognition composite score, crystallized cognition composite score) among the three groups (all p > 0.3). Compared with early middle-aged and middle-aged participants, older participants performed significantly worse on MoCA (p<0.001) and NIH working memory tests (p<0.001). Older participants had significantly larger BG-PVS volume (p<0.001) and CSO-PVS volume (p<0.001) than early middle-aged and middle-aged participants. In contrast, early middle-aged participants had significantly larger BG volume (p<0.0001) and CSO volume (p<0.0001) than middle-aged and older participants. Compared with early middle-aged and middle-aged participants, older participants had longer PSIQ self-rated sleep duration (p=0.017), and lower score for sleep medication (p=0.003). PSQI total score, sleep efficiency score, sleep quality score, sleep latency score, and time in bed were not significantly different among the three groups (all p > 0.1).

### 3.2 The relationship between PVS and brain functional connectivity (FC)

We first tested the associations between PVS VF and FC patterns in the whole sample. Figure 1 shows that higher PVS VF is associated with increased FC among various ROIs (Table 2). Specifically, BG-PVS VF was positively associated with FC between the right aMTG and a cluster in the right temporal region [cluster 1: 157 voxels, peak voxel MNI coordinates: +48, +14, -10; t(508)= 4.98], that included the right temporal pole (105 voxels) and the right insular cortex (38 voxels) (Figure 1a. r=0.21, p<0.0001). CSO-PVS VF was positively associated with FC of the left hippocampus and a cluster in the frontal regions [cluster 1: 145 voxels, peak voxel MNI coordinates: +48, +26, +02; t(508)= 4.70], including the right pars triangularis (105 voxels), the right pars opercularis (15 voxels), and the right frontal operculum cortex(10 voxels) (Figure 1b. r=0.21, p<0.0001).

**Figure1.**
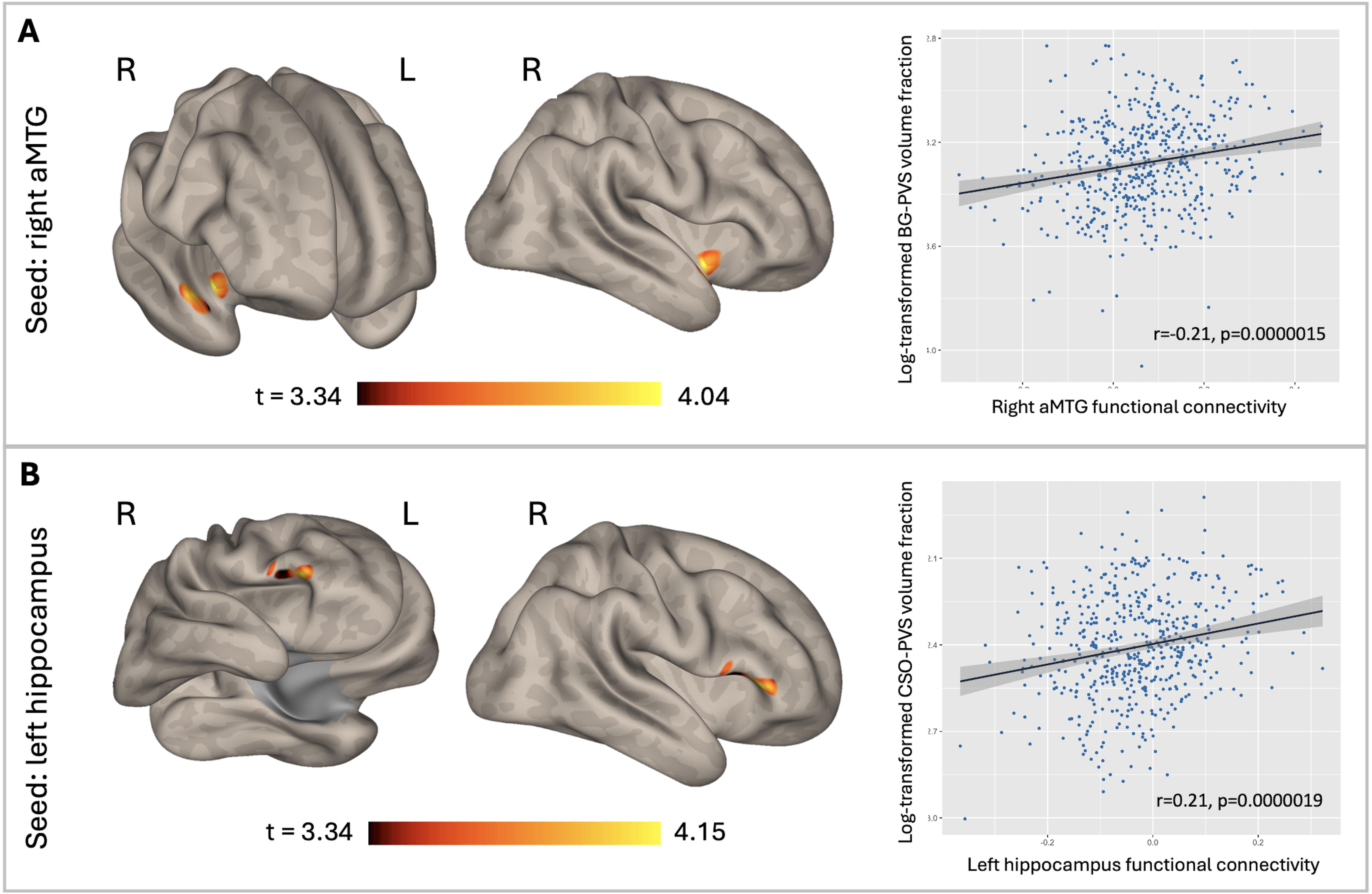
Brain functional connectivity (FC) related to PVS in HCP-Aging cohorts. **(A)** Brain activation map showing areas with BOLD response related to BG-PVS in an HCP-Aging cohorts. After adjustment for age, sex, and years of education, the functional connectivity between the right aMTG and right temporal cluster positively correlated with BG-PVS volume fraction(VF). **(B)** Brain activation map showing areas with BOLD response related to CSO-PVS in an HCP-Aging cohorts. After adjustment for age, sex, and years of education, the functional connectivity between the left hippocampus and right frontal cluster positively correlated with CSO-PVS VF. The color-bar represents the t-value for the statistical test. Significant ***p<0.0001.

**Table 2.**
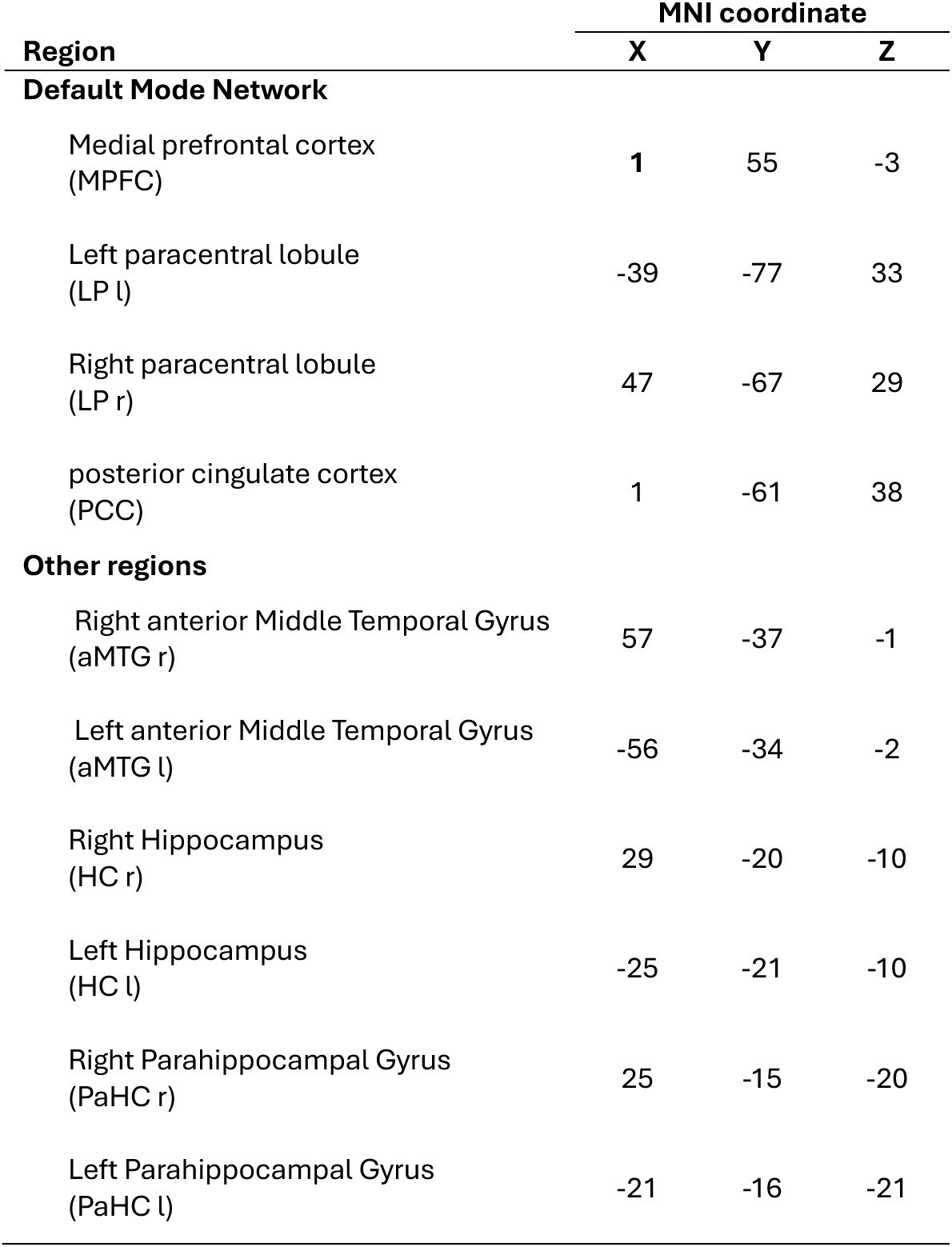
MNI coordinates of regions of interest (ROI)

| Region | MNI coordinate |  |  |
| --- | --- | --- | --- |
|  | X | Y | Z |
| **Default Mode Network** |  |  |  |
| Medial prefrontal cortex (MPFC) | 1 | 55 | -3 |
| Left paracentral lobule (LP l) | -39 | -77 | 33 |
| Right paracentral lobule (LP r) | 47 | -67 | 29 |
| posterior cingulate cortex (PCC) | 1 | -61 | 38 |
| **Other regions** |  |  |  |
| Right anterior Middle Temporal Gyrus (aMTG r) | 57 | -37 | -1 |
| Left anterior Middle Temporal Gyrus (aMTG l) | -56 | -34 | -2 |
| Right Hippocampus (HC r) | 29 | -20 | -10 |
| Left Hippocampus (HC l) | -25 | -21 | -10 |
| Right Parahippocampal Gyrus (PaHC r) | 25 | -15 | -20 |
| Left Parahippocampal Gyrus (PaHC l) | -21 | -16 | -21 |

### 3.3 Cognitive performance associated with FC

Our next question was whether the FC’s among the above mentioned regions (right aMTG-right temporal; left hippocampus-right frontal) were associated with cognition. FC between the aMTG and temporal regions, which was positively related to PVS in BG, was not significantly correlated with any cognition tests. However, in the case of the FC between the left hippocampus and frontal regions, which was positively related to PVS in CSO, a significant correlation was seen with cognition that varied across different age groups (Table 3).

**Table 3.**
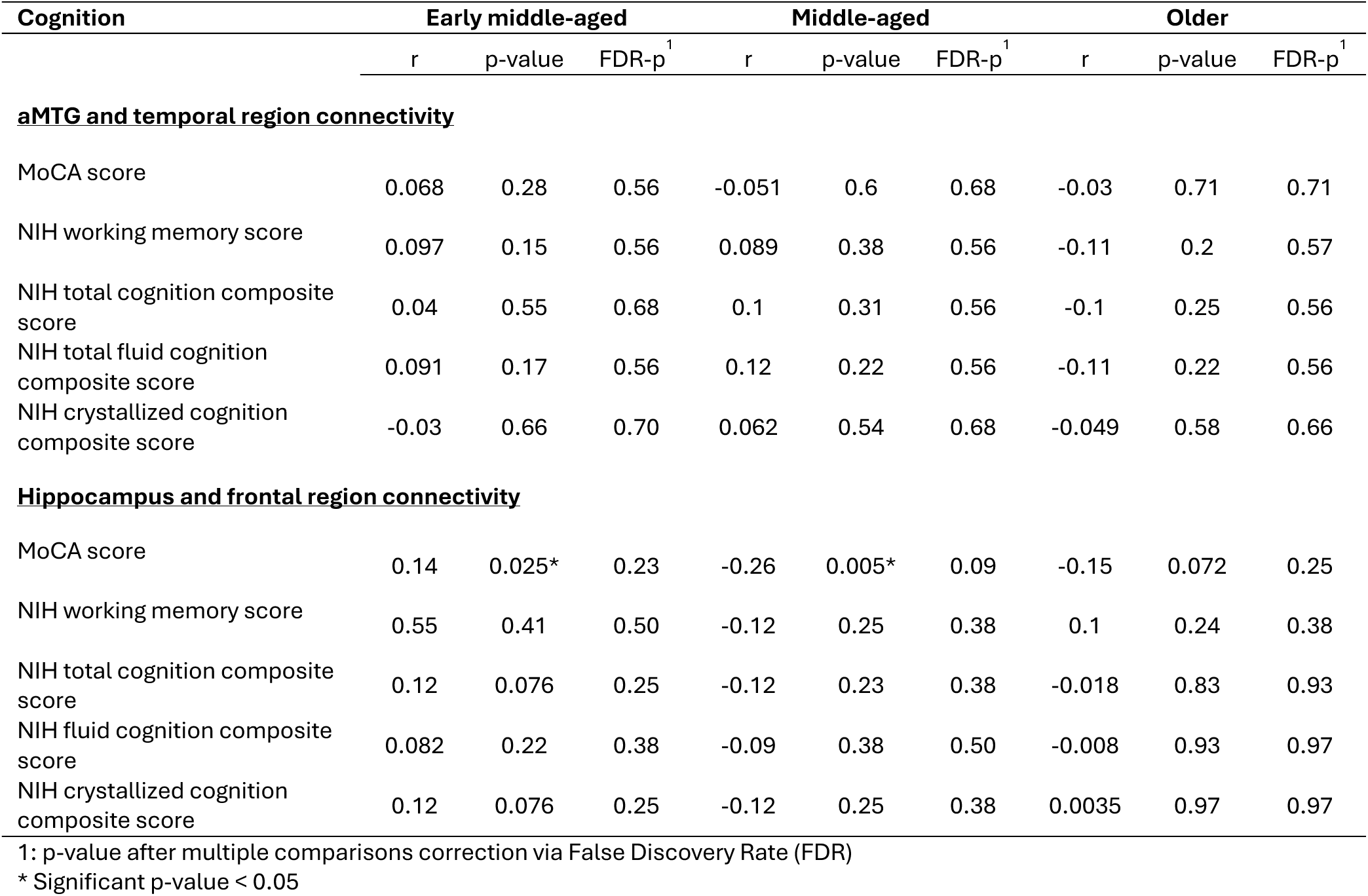
The correlation between PVS-related functional connectivity and cognitive performances.

| Cognition | Early middle-aged |  |  | Middle-aged |  |  | Older |  |  |
| --- | --- | --- | --- | --- | --- | --- | --- | --- | --- |
|  | r | p-value | FDR-p<sup>1</sup> | r | p-value | FDR-p<sup>1</sup> | r | p-value | FDR-p<sup>1</sup> |
| **aMTG and temporal region connectivity** |  |  |  |  |  |  |  |  |  |
| MoCA score | 0.068 | 0.28 | 0.56 | -0.051 | 0.6 | 0.68 | -0.03 | 0.71 | 0.71 |
| NIH working memory score | 0.097 | 0.15 | 0.56 | 0.089 | 0.38 | 0.56 | -0.11 | 0.2 | 0.57 |
| NIH total cognition composite score | 0.04 | 0.55 | 0.68 | 0.1 | 0.31 | 0.56 | -0.1 | 0.25 | 0.56 |
| NIH total fluid cognition composite score | 0.091 | 0.17 | 0.56 | 0.12 | 0.22 | 0.56 | -0.11 | 0.22 | 0.56 |
| NIH crystallized cognition composite score | -0.03 | 0.66 | 0.70 | 0.062 | 0.54 | 0.68 | -0.049 | 0.58 | 0.66 |
| **Hippocampus and frontal region connectivity** |  |  |  |  |  |  |  |  |  |
| MoCA score | 0.14 | 0.025* | 0.23 | -0.26 | 0.005* | 0.09 | -0.15 | 0.072 | 0.25 |
| NIH working memory score | 0.55 | 0.41 | 0.50 | -0.12 | 0.25 | 0.38 | 0.1 | 0.24 | 0.38 |
| NIH total cognition composite score | 0.12 | 0.076 | 0.25 | -0.12 | 0.23 | 0.38 | -0.018 | 0.83 | 0.93 |
| NIH fluid cognition composite score | 0.082 | 0.22 | 0.38 | -0.09 | 0.38 | 0.50 | -0.008 | 0.93 | 0.97 |
| NIH crystallized cognition composite score | 0.12 | 0.076 | 0.25 | -0.12 | 0.25 | 0.38 | 0.0035 | 0.97 | 0.97 |

1: p-value after multiple comparisons correction via False Discovery Rate (FDR)
\* Significant p-value < 0.05

Early middle-aged participants who had higher hippocampal FC showed increased MoCA score (r=0.14, p = 0.025). In contrast, the middle-aged participants who had higher hippocampal FC showed decreased MoCA score (r=-0.26, p = 0.005). However, statistical significance did not remain after multiple comparison correction. In addition, no correlation between PVS-related FC and cognition tests were found in the older group (all p>0.25). Upon further analysis of the link between PVS, functional connectivity, and cognition, hippocampal FC showed a mediating effect between PVS and MoCA score in the early middle-aged group (Figure 2A, average causal mediation effect (95% CI)= 0.393 (0.071, 0.84), p=0.013), but there was no statistically significant mediating effect in the middle-aged group (Figure 2B, average causal mediation effect (95% CI)= -0.488 (-1.133, 0.05), p=0.079).

**Figure 2.**
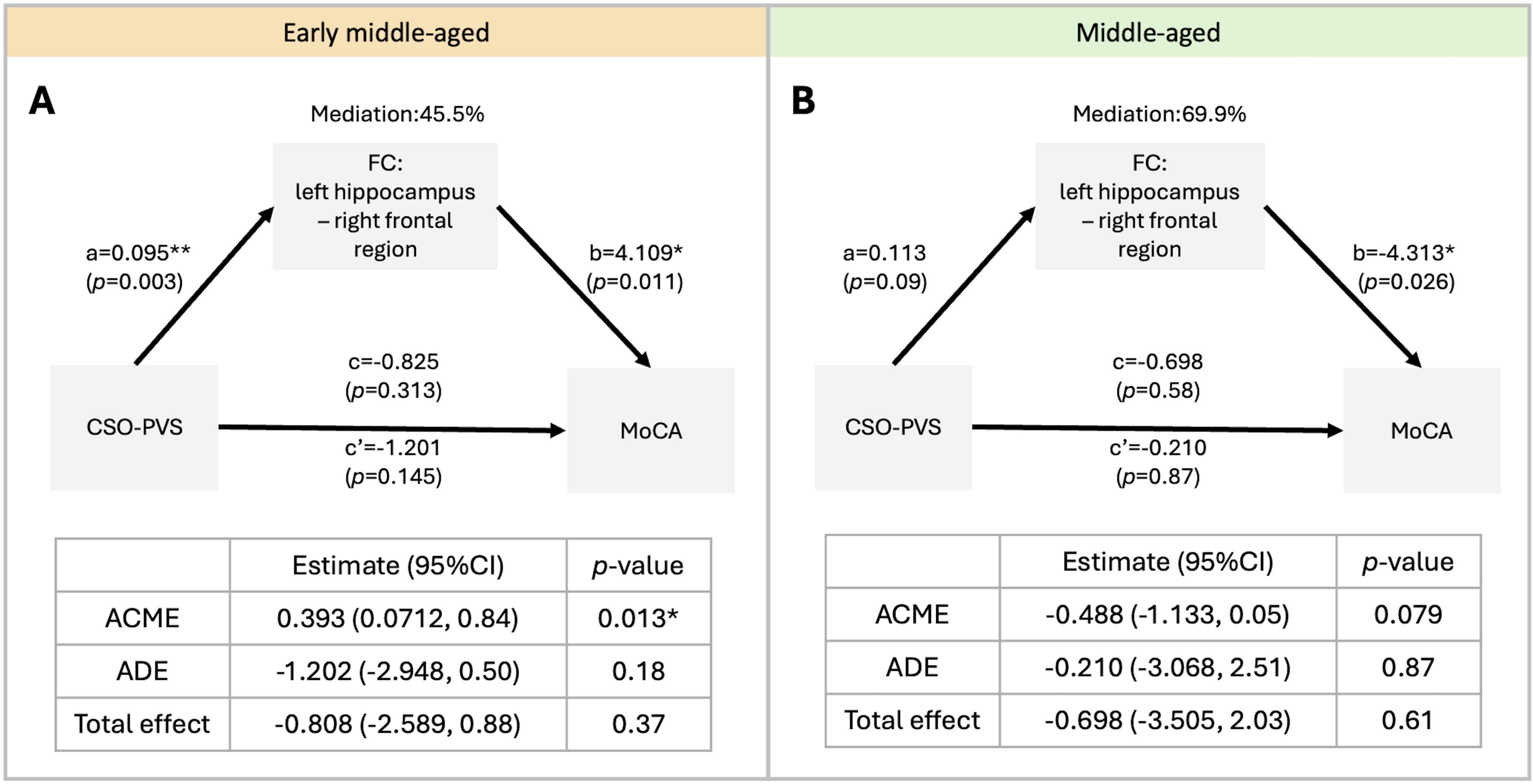
Demonstration of the mediation analysis of the CSO-PVS, FC and cognitive performance in the early middle-aged and middle-aged participants. **(A)** In early middle-aged population, there is mediating effect of hippocampus functional connectivity on the association between CSO-PVS volume fraction (VF) and the MoCA score. **(B)** In the older population, there is no mediating effect of aMTG functional connectivity on the association between CSO-PVS VF and MoCA score. Mediation (%) = (c-c’)/c. ACME: Average Causal Mediated Effect; ADE: Average Direct Effect. Significant *p<0.05.

### 3.4 The relationship between sleep and FC

Our next objective was to investigate whether sleep was associated with PVS-related FCs. These PVS-related FCs and PSQI sleep components did not show any significant correlation. (Table 4). To further investigate the effect of sleep quality on these PVS-related FCs, we grouped the participants based on the total PSQI score into a poor sleep group (score > 5) and a good sleep group (score <= 5). Right aMTG FC did not differ significantly between the two sleep groups (p=0.76). As well, left hippocampus FC did not differ significantly between poor and good sleep groups (Supplemental Figure 2A, p=0.93). In addition, the relationships between BG-PVS VF and right aMTG FC did not significantly differ between poor and good sleep groups (Supplemental Figure 2B). The relationships between CSO-PVS VF and left hippocampus FC also did not significantly differ between poor and good sleep groups (Supplemental Figure 2C).

**Table 4.** The correlation between functional connectivity and sleep measurements.

| Sleep measurements | Early middle-aged |  |  | Middle-aged |  |  | Older |  |  |
| --- | --- | --- | --- | --- | --- | --- | --- | --- | --- |
|  | r | p-value | FDR-p<sup>1</sup> | r | p-value | FDR-p<sup>1</sup> | r | p-value | FDR-p<sup>1</sup> |
| **<u>aMTG and temporal region connectivity</u>** |  |  |  |  |  |  |  |  |  |
| Total PSQI score | 0.11 | 0.077 | 0.64 | -0.061 | 0.53 | 0.91 | -0.067 | 0.42 | 0.68 |
| Sleep quality score | -0.027 | 0.67 | 0.96 | 0.016 | 0.87 | 0.94 | 0.046 | 0.58 | 0.71 |
| Sleep efficiency score | 0.11 | 0.082 | 0.4 | -0.18 | 0.052 | 0.34 | -0.09 | 0.27 | 0.68 |
| Sleep latency score | 0.09 | 0.15 | 0.45 | -0.007 | 0.94 | 0.94 | -0.074 | 0.37 | 0.68 |
| Sleep duration | -0.073 | 0.25 | 0.6 | 0.18 | 0.057 | 0.34 | 0.084 | 0.31 | 0.68 |
| Time in bed | 0.038 | 0.55 | 0.94 | 0.1 | 0.28 | 0.91 | -0.033 | 0.69 | 0.71 |
| **<u>Hippocampus and frontal region connectivity</u>** |  |  |  |  |  |  |  |  |  |
| Total PSQI score | -0.017 | 0.78 | 0.96 | 0.033 | 0.73 | 0.94 | 0.064 | 0.44 | 0.68 |
| Sleep quality score | 0.007 | 0.91 | 0.96 | 0.064 | 0.51 | 0.91 | 0.11 | 0.19 | 0.68 |
| Sleep efficiency score | -0.003 | 0.96 | 0.96 | -0.011 | 0.91 | 0.94 | 0.062 | 0.45 | 0.68 |
| Sleep latency score | -0.045 | 0.48 | 0.94 | -0.073 | 0.45 | 0.91 | -0.085 | 0.3 | 0.68 |
| Sleep duration | -0.006 | 0.93 | 0.96 | -0.035 | 0.71 | 0.94 | -0.037 | 0.65 | 0.71 |
| Time in bed | 0.1 | 0.1 | 0.4 | -0.085 | 0.38 | 0.91 | 0.03 | 0.71 | 0.71 |

1: p-value after multiple comparisons correction via False Discovery Rate (FDR)

### 3.5 The relationship between PVS, FC, and sleep

Despite the lack of a statistically significant correlation between sleep and FC in the overall sample in this study, we explored if these associations differed over age groups. The correlation between PVS and sleep parameters varied across different age groups according to our previous research^21^, and we found a similar pattern in this study (Supplemental Table 1). In the early middle-aged group, mediation analysis revealed a significant mediation effect of time in bed on right aMTG FC through BG-PVS VF (Figure 3A, average causal mediation effect (95% CI) = 0.0011 (0.000089, 0.00), p=0.027). In addition, there was a marginally significant mediation effect of time in bed on hippocampus FC through CSO-PVS VF (Figure 4B, average causal mediation effect (95% CI) = 0.0006 (-1.19e-05, 0.00), p=0.058). Furthermore, in the older group, the result indicated a significant mediation effect of sleep quality scores on right aMTG FC through BG-PVS VF (Figure 3C, average causal mediation effect (95% CI) = -0.0109 (-0.025, 0.00), p=0.009).

**Figure 3.**
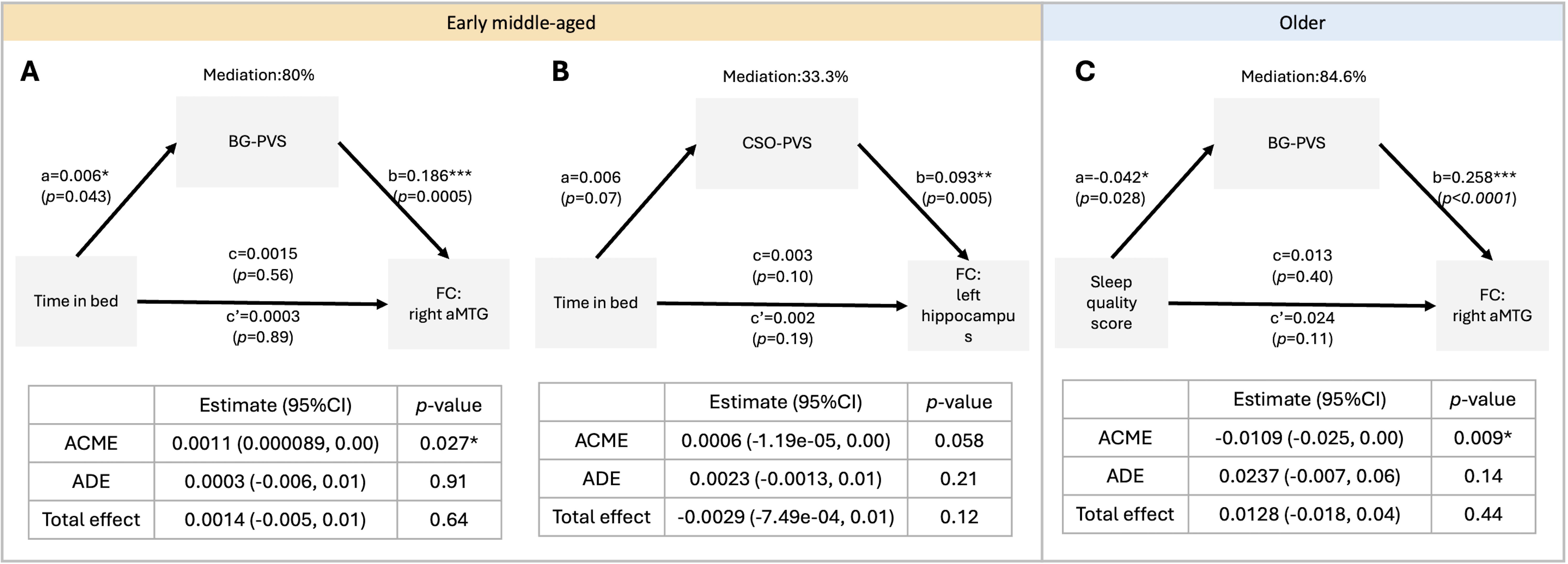
Demonstration of the mediation analysis of the PVS, time in bed/sleep quality and functional connectivity in the early middle-aged and older participants. **(A)** In the early middle-aged cohort, there is mediating effect of BG-PVS volume fraction (VF) on the association between time in bed and the aMTG functional connectivity; however, **(B)** there is no mediating effect of CSO-PVS VF on the association between time in bed and the hippocampus functional connectivity. **(C)** In the older cohort, there is mediating effect of BG-PVS VF on the association between time in bed and the aMTG functional connectivity. Mediation (%) = (c-c’)/c. ACME: Average Causal Mediated Effect; ADE: Average Direct Effect. Significant *p<0.05.

## 4 Discussion

This study is the first to explore the relationship between PVS and FC. Our results indicate that people who have larger PVS in the BG showed higher FC between right aMTG and right temporal pole and insular cortex. People who had larger CSO-PVS showed higher FC between left hippocampus and right frontal regions: right pars triangularis, right pars opercularis and right frontal operculum cortex. In addition, the relationship between PVS, FC and cognition varied across the different age groups. In early middle-aged individuals (age: 36-55 years old), the connection between PVS and MoCA scores is mediated by PVS associations on FC of the hippocampus. Early middle-aged individuals who had larger CSO-PVS VF showed higher hippocampus FC, which was associated with better cognitive performance. However, we did not find any correlation between these FC and cognition in middle-aged group (age: 55-65 years old) and older group (age above 55 years old).

### 4.1 PVS and FC

The mechanism between PVS and FC is still unclear. However, the link between PVS and FC may contribute to the integration of vascular and neural processes that are crucial for cognitive function. One potential link could be that the function of the PVS affects interstitial fluid dynamics, which in turn affects the delivery and clearance of vasoactive substances that regulate blood flow, ultimately affecting the BOLD signal^36^. Previous research found that vasomotor activity is associated with the BOLD signal that reflects baseline oscillations in cerebral blood flow and oxygenation ^10^. Vasomotor activity is a physiological mechanism that regulates blood vessel diameter (vasodilation and vasoconstriction) by the smooth muscle in the vessel walls. Rhythmic changes in vessel diameter caused by vasomotion regulate local CBF and cerebral blood volume (CBV), thereby influencing the magnitude and dynamics of the BOLD response to neuronal activation^10^. Vasomotion during different sleep stages regulates CSF flow through the PVS, supporting waste clearance^37^. Impaired vasomotion patterns were observed in conditions such as AD, stroke, and hypertension^38–40^, highlighting its critical role in glymphatic function. Previous studies have pointed out that changes in the PVS structure can affect the contraction and relaxation of blood vessels by changing the mechanical properties of blood vessels and disrupting neurovascular signal conduction^10,36^. When the PVS is enlarged or dysfunctional, it may change the mechanical properties of peripheral blood vessels^41^, affecting the transmural pressure of the vessel walls, thereby changing the diastolic tension and pattern of the vessel. In addition, the PVS is a key exchange channel for fluid and signaling molecules around the cerebral blood vessels^1^. Change to these pathways may impact the regulation of vascular tone and affect the vasomotion function. The glymphatic system facilitates the exchange of ions and metabolites between blood vessels and nervous tissue. Any imbalance will affect the smooth muscle cells of the blood vessel wall, thereby changing the blood vessel’s ability to relax and contract^42,43^. PVS dysfunction leads to impaired clearance of metabolic waste products, and accumulated waste products trigger inflammation^1^, leading to changes in vascular tone and reactivity^44^. In AD, changes in PVS may lead to impaired vasomotion due to amyloid-β deposition, thereby affecting the elasticity and function of blood vessels^39^. PVS is also involved in neurovascular signaling pathways, and these vasomotor activities in turn affect local blood flow and oxygenation levels, thereby modulating the BOLD signal used in fMRI to infer neural activity^10,36^.

In addition, a previous study reported that synchronized neuronal oscillations, particularly during sleep, create ionic waves that help drive CSF movement, enhancing the clearance of waste, including amyloid-β proteins^45^. Disruptions to these neuronal dynamics impair glymphatic function, reducing the efficiency of waste clearance ^1^, while enhanced neuronal activity can improve the waste clearance^45^. Enlarged PVS, which increases CSF flow, could potentially impact brain FC, particularly by enhancing the glymphatic system’s efficiency in clearing metabolic waste and promoting optimal interstitial fluid dynamics, despite the enlarged PVS is always associated with pathology, which makes it not a positive thing. Furthermore, our results showed that people who had better sleep quality or spent more time in bed had larger PVS and higher FC. Previous research that evaluated associations of sleep with PVS volume hypothesized that there may be a compensatory mechanism in which fluid accumulates because perivascular/glymphatic system dysfunction results in less sleep^46–48^. Another animal study has shown that vascular dynamics during different sleep stages influence CSF flow and solute transport through the perivascular space. Slow-wave sleep (NREM) is associated with slow, large-amplitude oscillations in arteries that enhance the movement of fluids and solutes, while REM sleep causes vasodilation and reduced flow. These sleep-related vascular changes can significantly improve brain waste clearance^37^. The above evidence suggested that the enlargement or dysfunction of the PVS may affect the vascular dynamics and neural activity of the brain through multiple mechanisms, thereby affecting the FC.

On the other hand, a recent fMRI study reported that large cortical BOLD signals are accompanied by strong CSF movement, possibly indicating active cortical glymphatic function^49^. During sleep, large fluctuations in the global BOLD signal are also accompanied by CSF influx^49^. Furthermore, some studies suggest that enlargement of the PVS may be due to CSF flow. PVS are involved in the drainage of brain interstitial fluid, and PVS enlargement may occur to compensate for the increase in CSF flow, especially in situations where CSF production is increased or other clearance pathways are impaired, forcing the PVS to process more CSF^50,51^. The compensatory mechanism hypothesis proposes that the enlarged PVS is able to accommodate increased fluid flow due to changes in interstitial hydrodynamics, although this hypothesis is still under discussion. In addition, changes in PVS affect the brain’s ability to manage metabolic waste, and increased CSF flow may affect the brain’s FC by improving the clearance efficiency of metabolic waste by the glymphatic system and promoting optimal interstitial fluid dynamics. Previous studies have shown that large waves of CSF inflow are closely related to changes in the BOLD signal^52^. Although there is no direct evidence that increased CSF is associated with enhanced FC in the brain, some hypotheses support this possibility: (1) Increased CSF flow can enhance waste clearance in the brain ^53^, reduce neuroinflammation, and promote better functional connection, as efficient waste removal during periods of high metabolic activity is critical to maintaining a clean and functional neural environment; (2) CSF plays a key role in distributing nutrients to different brain regions^54^, and enhanced CSF dynamics can Improve nutrient delivery and support synaptic activity and communication of neural circuits, thereby enhancing functional connectivity; (3) Some studies have shown that CSF affects important brain areas related to memory, visual construction, verbal fluency, and cognition, enhancing CSF may alter FC patterns in these regions^55,56^, slowing down neurodegenerative processes and improving cognitive function. Taken together, enlargement of PVS may affect the brain’s FC and overall cognitive health through multiple mechanisms.

In our study, enlarged PVS was associated with higher FC. Our results suggest that increased PVS that may promote increased brain FC may be as a compensatory mechanism to maintain cognitive function to prevent the potential impact of enlarged BG-PVS on cognitive function in healthy older adults. One study showed that healthy older adults with enlarged BG-PVS showed higher connectivity coupling between retrosplenial temporal network and cingulo-parietal network compared to those without enlarged PVS^57^. Another longitudinal study showed that enlarged PVS burden in the BG was positively associated with declines in language, information processing, executive functions, and contextual memory^44^. These findings show that alterations in microvasculature are associated with changes in FC.

Although our results indicated that sleep quality does not significantly affect the relationship between PVS VF and FC, after investigating deeper into the relationship between PVS, sleep and FC, the results indicated that early middle-aged people with longer time in bed had a larger BG-PVS VF that was associated with higher levels of functional connectivity on the right aMTG. On the other hand, older people with better sleep quality had a larger BG-PVS VF that was associated with higher levels of FC on the right aMTG. However, we did not find any correlation between PVS and sleep in the middle-aged group.

### 4.2 FC between aMTG and temporal region

We found that individuals with larger BG-PVS showed higher FC in the right aMTG and a cluster in right temporal regions (temporal pole and insula cortex), which are involved in language and memory-related processing. The MTG is involved in language and semantic memory processing, visual perception, and multimodal sensory integration. Recent advancements in tractography-based parcellation have pointed out that the aMTG is primarily responsible for sound recognition and semantic retrieval ^58^. Connectivity Between MTG, temporal pole, and Insular cortex are interconnected through several white matter tracts. The MTG and temporal pole are connected via the uncinate fasciculus, a major white matter tract linking the anterior temporal lobe to the prefrontal cortex and insular regions^59^. The insula also has connections with both the MTG and temporal pole through the inferior fronto-occipital fasciculus and the middle longitudinal fasciculus^59^. Resting-State functional MRI studies show that the MTG, temporal pole, and insular cortex often co-activate during tasks for the integration of sensory information with emotional and memory processes, supporting complex cognitive functions like empathy, decision-making, and language comprehension^60^. A meta-analysis has also highlighted the MTG’s involvement in the DMN^61^. In addition, there is a strong anatomical connectivity between aMTG and the medial temporal lobe, encompassing structures like the hippocampus, lingual gyrus, and fusiform gyrus^58^. Therefore, changes in aMTG-temporal region connectivity may represent an alteration in language and memory processing.

### 4.3 FC between hippocampus and frontal region

We found that individuals with larger CSO-PVS showed higher FC between left hippocampus and right frontal region (right pars triangularis, right pars opercularis, and the right frontal operculum cortex), areas critical for language and memory processing. The hippocampus is crucial for memory formation, spatial navigation, and the consolidation of information from short-term to long-term memory^33^. It is part of the limbic system and plays a significant role in emotional regulation and learning^33^. The frontal operculum encompasses the pars triangularis and pars opercularis, which are subregions of the inferior frontal gyrus (IFG)^62^. The pars triangularis and pars opercularis are involved in language processing and speech production^63^. Functional connectivity between the hippocampus and IFG regions is often associated with the integration of memory and language^64^. This connectivity is crucial during tasks that require verbal memory recall or when processing language in the context of past experiences^64,65^. Previous research has shown decreased FC between the pars opercularis and the left hippocampus after total sleep deprivation is linked to phonological loop impairment^66^. However, there was no direct correlation observed between any sleep parameters and FCs in this study.

### 4.4 Limitations

This study still needs to note some limitations. Among them, the fMRI ROIs are located in gray matter, further away from the studied PVS presentation regions. The integration of interpretations between white matter structural changes and gray matter neural activity is a major challenge of this study, and further tracer studies are required. Additionally, the relationship between PVS and FC was only observed at one time point. Therefore, future research needs to plan longitudinal studies to accurately investigate the long-term impact of alterations in PVS on FC changes. Furthermore, the majority of the participants in the study were white or African American, so these findings might not apply to other populations.

### 4.5 Conclusion

In conclusion, our results demonstrated that BG-PVS VF was positively associated with FC of the right aMTG and a cluster in temporal regions; CSO-PVS VF was positively associated with FC of the left hippocampus and a cluster in frontal IFG regions. In addition, FC in the studied seed regions play a critical role in mediating the relationship between PVS and cognition ability. Furthermore, despite sleep not directly affecting the studied FCs, PVS in BG and CSO mediated the interaction between time in bed and FCs in the early middle-aged population; while in the older population, only BG-PVS mediated the interaction between sleep quality and right aMTG FCs. This study could be used as a foundation for future research relating PVS, FC, and sleep behavior. The findings of this study provide a greater understanding of how PVS alternation influences brain FC in older, cognitively healthy individuals, and open a new area of PVS research (structure-to-function) to investigation.

## Acknowledgments

The research reported in this publication was supported by the National Institute of Mental Health, and National Institute on Aging of the NIH under Award Numbers RF1MH123223, and R01AG070825. The content is solely the responsibility of the authors and does not necessarily represent the official views of the NIH.

## HCP-Aging

Dataset used in this publication was supported by the National Institute on Aging of the National Institutes of Health under Award Number U01AG052564. The content is solely the responsibility of the authors and does not necessarily represent the official views of the National Institutes of Health.

## Author contributions

N.S., J.C., W.M., and J.C. conceived the research study. N.S., and J.C. analyzed and interpreted the data. N.S. wrote the manuscript. J.C., W.M. and J.C. reviewed the manuscript critically. All authors edited and revised the manuscript and approved final submission.

## References

1. Barisano G, Lynch KM, Sibilia F, et al. Imaging perivascular space structure and function using brain MRI. Neuroimage 2022; 257: 119329.

2. Yue Y, Zhang X, Lv W, et al. Interplay between the glymphatic system and neurotoxic proteins in Parkinson’s disease and related disorders: current knowledge and future directions. Neural Regeneration Research 2024; 19: 1973–1980.

3. Shokri-Kojori E, Wang GJ, Wiers CE, et al. β-Amyloid accumulation in the human brain after one night of sleep deprivation. Proc Natl Acad Sci U S A 2018; 115: 4483– 4488.

4. Semyachkina-Glushkovskaya O, Postnov D, Penzel T, et al. Sleep as a novel biomarker and a promising therapeutic target for cerebral small vessel disease: A review focusing on alzheimer’s disease and the blood-brain barrier. International Journal of Molecular Sciences 2020; 21: 1–15.

5. Xie L, Kang H, Xu Q, et al. Sleep drives metabolite clearance from the adult brain. Science (1979) 2013; 342: 373–7.

6. Sepehrband F, Barisano G, Sheikh-Bahaei N, et al. Volumetric distribution of perivascular space in relation to mild cognitive impairment. Neurobiol Aging 2021; 99: 28–43.

7. Shih N, Barisano G, Lincoln KD, et al. Effects of sleep on brain perivascular space in a cognitively healthy population. Sleep Med 2023; 111: 170–179.

8. Lysen TS, Zonneveld HI, Muetzel RL, et al. Sleep and resting-state functional magnetic resonance imaging connectivity in middle-aged adults and the elderly: A population-based study. J Sleep Res 2020; 29: 1–10.

9. Andrews-Hanna JR. The brain’s default network and its adaptive role in internal mentation. Neuroscientist 2012; 18: 251–270.

10. Hillman EMC. Coupling mechanism and significance of the BOLD signal: A status report. Annual Review of Neuroscience 2014; 37: 161–181.

11. Heeger DJ, Ress D. What does fMRI tell us about neuronal activity? Nature Reviews Neuroscience 2002; 3: 142–151.

12. Ingala S, Tomassen J, Collij LE, et al. Amyloid-driven disruption of default mode network connectivity in cognitively healthy individuals. Brain Commun; 3. Epub ahead of print 2021. DOI: 10.1093/braincomms/fcab201.

13. Palmqvist S, Schöll M, Strandberg O, et al. Earliest accumulation of β-amyloid occurs within the default-mode network and concurrently affects brain connectivity. Nat Commun; 8. Epub ahead of print 2017. DOI: 10.1038/s41467-017-01150-x.

14. Zhang Y, Dai C, Shao Y, et al. Decreased Functional Connectivity in the Reward Network and Its Relationship With Negative Emotional Experience After Total Sleep Deprivation. Front Neurol 2021; 12: 1–10.

15. Hehr A, Huntley ED, Marusak HA. Getting a Good Night’s Sleep: Associations Between Sleep Duration and Parent-Reported Sleep Quality on Default Mode Network Connectivity in Youth. Journal of Adolescent Health 2023; 72: 933–942.

16. Verweij IM, Romeijn N, Smit DJA, et al. Sleep deprivation leads to a loss of functional connectivity in frontal brain regions. BMC Neurosci; 15. Epub ahead of print 19 July 2014. DOI: 10.1186/1471-2202-15-88.

17. Dai C, Zhang Y, Cai X, et al. Effects of Sleep Deprivation on Working Memory: Change in Functional Connectivity Between the Dorsal Attention, Default Mode, and Fronto-Parietal Networks. Front Hum Neurosci; 14. Epub ahead of print 12 October 2020. DOI: 10.3389/fnhum.2020.00360.

18. Ward AM, McLaren DG, Schultz AP, et al. Daytime sleepiness is associated with decreased default mode network connectivity in both young and cognitively intact elderly subjects. Sleep 2013; 36: 1609–1615.

19. Chou KH, Kuo CY, Liang CS, et al. Shared patterns of brain functional connectivity for the comorbidity between migraine and insomnia. Biomedicines 2021; 9: 1–17.

20. Bookheimer SY, Salat DH, Terpstra M, et al. The Lifespan Human Connectome Project in Aging: An overview. Neuroimage 2019; 185: 335–348.

21. Shih N, Barisano G, Lincoln KD, et al. Effects of sleep on brain perivascular space in a cognitively healthy population. Sleep Med 2023; 111: 170–179.

22. Lachman S, Boekholdt SM, Luben RN, et al. Impact of physical activity on the risk of cardiovascular disease in middle-aged and older adults: EPIC Norfolk prospective population study. Eur J Prev Cardiol 2018; 25: 200–208.

23. Hafkemeijer A, Altmann-Schneider I, de Craen AJM, et al. Associations between age and gray matter volume in anatomical brain networks in middle-aged to older adults. Aging Cell 2014; 13: 1068–1074.

24. Gale SC, Peters JA, Murry JS, et al. Injury patterns and outcomes in late middle age (55–65): The intersecting comorbidity with high-risk activity – A retrospective cohort study. Annals of Medicine and Surgery 2018; 27: 22–25.

25. Brown WR, Moody DM, Thore CR, et al. Vascular dementia in leukoaraiosis may be a consequence of capillary loss not only in the lesions, but in normal-appearing white matter and cortex as well. J Neurol Sci 2007; 257: 62–66.

26. Glasser MF, Sotiropoulos SN, Wilson JA, et al. The minimal preprocessing pipelines for the Human Connectome Project. Neuroimage 2013; 80: 105–124.

27. Whitfield-Gabrieli S, Nieto-Castanon A. Conn: A Functional Connectivity Toolbox for Correlated and Anticorrelated Brain Networks. Brain Connect 2012; 2: 125–141.

28. Sepehrband F, Barisano G, Sheikh-Bahaei N, et al. Image processing approaches to enhance perivascular space visibility and quantification using MRI. Sci Rep 2019; 9: 1–12.

29. Cabeen RP, Laidlaw DH, Toga AW. Quantitative Imaging Toolkit : Software for Interactive 3D Visualization, Data Exploration, and Computational Analysis of Neuroimaging Datasets. ISMRM-ESMRMB Abstracts 2018; 12–14.

30. Penny WD, FKJ, AJT, KSJ, & NTE. Statistical parametric mapping: the analysis of functional brain images. Elsevier.

31. Friston KJ, Williams S, Howard R, et al. Movement-related effects in fMRI time-series. Magn Reson Med 1996; 35: 346–355.

32. Nieto-Castanon A. Handbook of functional connectivity Magnetic Resonance Imaging methods in CONN. Hilbert Press, 2020. Epub ahead of print 4 February 2020. DOI: 10.56441/hilbertpress.2207.6598.

33. Pronier É, Morici JF, Girardeau G. The role of the hippocampus in the consolidation of emotional memories during sleep. Trends in Neurosciences 2023; 46: 912–925.

34. Sepehrband F, Barisano G, Sheikh-Bahaei N, et al. Volumetric distribution of perivascular space in relation to mild cognitive impairment. Neurobiol Aging 2021; 99: 28–43.

35. Buysse DJ, Reynolds CF, Monk TH, et al. The Pittsburgh Sleep Quality Index: a new instrument for psychiatric practice and research. Psychiatry Res 1989; 28: 193–213.

36. Drew PJ. Vascular and neural basis of the BOLD signal. Current Opinion in Neurobiology 2019; 58: 61–69.

37. Bojarskaite L, Vallet A, Bjørnstad DM, et al. Sleep cycle-dependent vascular dynamics in male mice and the predicted effects on perivascular cerebrospinal fluid flow and solute transport. Nat Commun; 14. Epub ahead of print 1 December 2023. DOI: 10.1038/s41467-023-36643-5.

38. Santisteban MM, Iadecola C, Carnevale D. Hypertension, Neurovascular Dysfunction, and Cognitive Impairment. Hypertension 2023; 80: 22–34.

39. Fisher RA, Miners JS, Love S. Pathological changes within the cerebral vasculature in Alzheimer’s disease: New perspectives. Brain Pathology; 32. Epub ahead of print 1 November 2022. DOI: 10.1111/bpa.13061.

40. Liu M, Zhang X, Wang B, et al. Functional status of microvascular vasomotion is impaired in spontaneously hypertensive rat. Sci Rep; 7. Epub ahead of print 1 December 2017. DOI: 10.1038/s41598-017-17013-w.

41. Ramaswamy S, Khasiyev F, Gutierrez J. Brain Enlarged Perivascular Spaces as Imaging Biomarkers of Cerebrovascular Disease: A Clinical Narrative Review. Journal of the American Heart Association; 11. Epub ahead of print 20 December 2022. DOI: 10.1161/JAHA.122.026601.

42. Brozovich F V., Nicholson CJ, Degen C V., et al. Mechanisms of vascular smooth muscle contraction and the basis for pharmacologic treatment of smooth muscle disorders. Pharmacological Reviews 2016; 68: 476–532.

43. Touyz RM, Alves-Lopes R, Rios FJ, et al. Vascular smooth muscle contraction in hypertension. Cardiovascular Research 2018; 114: 529–539.

44. Bown CW, Khan OA, Liu D, et al. Enlarged perivascular space burden associations with arterial stiffness and cognition. Neurobiol Aging 2023; 124: 85–97.

45. Jiang-Xie L-F, Drieu A, Bhasiin K, et al. Neuronal dynamics direct cerebrospinal fluid perfusion and brain clearance. Nature 2024; 627: 157–164.

46. Aribisala BS, Riha RL, Valdes Hernandez M, et al. Sleep and brain morphological changes in the eighth decade of life. Sleep Med 2020; 65: 152–158.

47. Ramirez J, Holmes MF, Berezuk C, et al. MRI-visible perivascular space volumes, sleep duration and daytime dysfunction in adults with cerebrovascular disease. Sleep Med 2021; 83: 83–88.

48. Lysen TS, Yilmaz P, Dubost F, et al. Sleep and perivascular spaces in the middle-aged and elderly population. J Sleep Res 2021; 1–9.

49. Han F, Chen J, Belkin-Rosen A, et al. Reduced coupling between cerebrospinal fluid flow and global brain activity is linked to Alzheimer disease–related pathology. PLoS Biol; 19. Epub ahead of print 1 June 2021. DOI: 10.1371/journal.pbio.3001233.

50. Richmond SB, Seidler RD, Iliff JJ, et al. Dynamic changes in perivascular space morphology predict signs of spaceflight-associated neuro-ocular syndrome in bed rest. NPJ Microgravity; 10. Epub ahead of print 1 December 2024. DOI: 10.1038/s41526-024-00368-6.

51. Garic D, McKinstry RC, Rutsohn J, et al. Enlarged Perivascular Spaces in Infancy and Autism Diagnosis, Cerebrospinal Fluid Volume, and Later Sleep Problems. JAMA Netw Open 2023; E2348341.

52. Fultz NE, Bonmassar G, Setsompop K, et al. Coupled electrophysiological, hemodynamic, and cerebrospinal fluid oscillations in human sleep. Science (1979) 2019; 366: 628–631.

53. Lysen TS, Yilmaz P, Dubost F, et al. Sleep and perivascular spaces in the middle-aged and elderly population. J Sleep Res 2021; 1–9.

54. Spector R, Robert Snodgrass S, Johanson CE. A balanced view of the cerebrospinal fluid composition and functions: Focus on adult humans. Experimental Neurology 2015; 273: 57–68.

55. Attier-Zmudka J, Sérot JM, Valluy J, et al. Decreased cerebrospinal fluid flow is associated with cognitive deficit in elderly patients. Front Aging Neurosci; 11. Epub ahead of print 2019. DOI: 10.3389/fnagi.2019.00087.

56. Hazan J, Wing M, Liu KY, et al. Clinical utility of cerebrospinal fluid biomarkers in the evaluation of cognitive impairment: A systematic review and meta-analysis. *Journal of Neurology*, Neurosurgery and Psychiatry 2022; 94: 113–120.

57. Zhang H, Cao P, Mak HKF, et al. The structural–functional-connectivity coupling of the aging brain. Geroscience 2024; 46: 3875–3887.

58. Xu J, Wang J, Fan L, et al. Tractography-based Parcellation of the Human Middle Temporal Gyrus. Nature Publishing Group 2015; 1–13.

59. Catani M, Thiebaut de Schotten M. A diffusion tensor imaging tractography atlas for virtual in vivo dissections. Cortex 2008; 44: 1105–1132.

60. Wang J, Yang Z, Klugah-Brown B, et al. The critical mediating roles of the middle temporal gyrus and ventrolateral prefrontal cortex in the dynamic processing of interpersonal emotion regulation. Neuroimage 2024; 120789.

61. Binder JR, Desai RH, Graves WW, et al. Where Is the Semantic System ? A Critical Review and Meta-Analysis of 120 Functional Neuroimaging Studies. Epub ahead of print 2009. DOI: 10.1093/cercor/bhp055.

62. John H. Martin. Neuroanatomy Text and Atlas, Fourth Edition. McGraw Hill Professional.

63. Eggermont JJ. Chapter 10 - Brain networks involved in deviance and novelty detection: Are they sensory modality specific? In: Eggermont JJ (ed) Brain Responses to Auditory Mismatch and Novelty Detection. Academic Press, pp. 315–343.

64. Raud L, Sneve MH, Vidal-Piñeiro D, et al. Hippocampal-cortical functional connectivity during memory encoding and retrieval. Neuroimage; 279. Epub ahead of print 1 October 2023. DOI: 10.1016/j.neuroimage.2023.120309.

65. Preston AR, Eichenbaum H. Interplay of hippocampus and prefrontal cortex in memory. Current Biology; 23. Epub ahead of print 9 September 2013. DOI: 10.1016/j.cub.2013.05.041.

66. Wang L, Wu H, Dai C, et al. Dynamic hippocampal functional connectivity responses to varying working memory loads following total sleep deprivation. J Sleep Res; 32. Epub ahead of print 1 June 2023. DOI: 10.1111/jsr.13797.

