## Supplementary for "Perivascular space mediated the interaction between sleep, and brain functional connectivity in the healthy aging population"

### Supplementary Table and figure

**Supplementary Table 1.** Multivariable linear regression analysis was conducted to investigate the relationship between the PVS volume fraction (VF) in basal ganglia (BG) and centrum semiovale (CSO) and sleep measurements across early middle-aged, middle-aged, and older groups.

| Dependent variable | Predictor | Early middle-aged | | | Middle-aged | | | Older | | |
|---|---|---|---|---|---|---|---|---|---|---|
| | | Coefficient beta | p-value | FDR-p[1] | Coefficient beta | p-value | FDR-p[1] | Coefficient beta | p-value | FDR-p[1] |
| BG-PVS VF | Total PSQI score | -0.0005 | 0.894 | 0.95 | 0.0005 | 0.907 | 0.95 | -0.0034 | 0.436 | 0.88 |
| | Sleep efficiency score | -0.0004 | 0.974 | 0.97 | -0.0016 | 0.921 | 0.97 | -0.023 | 0.108 | 0.65 |
| | Sleep quality score | 0.0046 | 0.757 | 0.97 | -0.0127 | 0.514 | 0.93 | -0.0429 | 0.025* | 0.31 |
| | Sleep latency score | -0.0081 | 0.494 | 0.88 | -0.0041 | 0.808 | 0.95 | 0.0098 | 0.5 | 0.88 |
| | Sleep duration | 0.0062 | 0.641 | 0.88 | 0.0061 | 0.543 | 0.88 | 0.0151 | 0.164 | 0.88 |
| | Time in bed | 0.0061 | 0.041* | 0.73 | -0.0073 | 0.152 | 0.88 | 0.0004 | 0.946 | 0.95 |
| CSO-PVS VF | Total PSQI score | -0.0039 | 0.338 | 0.93 | 0.0031 | 0.569 | 0.93 | 0.0027 | 0.560 | 0.93 |
| | Sleep efficiency score | 0.0078 | 0.610 | 0.88 | -0.0106 | 0.564 | 0.88 | -0.0144 | 0.362 | 0.88 |
| | Sleep quality score | -0.0066 | 0.683 | 0.88 | 0.0228 | 0.301 | 0.88 | 0.0167 | 0.427 | 0.88 |
| | Sleep latency score | -0.0114 | 0.378 | 0.93 | 0.0030 | 0.873 | 0.97 | -0.0092 | 0.563 | 0.93 |
| | Sleep duration | -0.0005 | 0.963 | 0.97 | -0.0064 | 0.671 | 0.97 | -0.0136 | 0.245 | 0.88 |
| | Time in bed | 0.0068 | 0.035* | 0.31 | -0.0078 | 0.174 | 0.78 | 0.00123 | 0.814 | 0.97 |

BG: Basal ganglia; CSO: Centrum semiovale; PVS: Perivascular space; VF: Volume fraction

1: p-value after multiple comparisons correction via False Discovery Rate (FDR)

* Significant p-value < 0.05

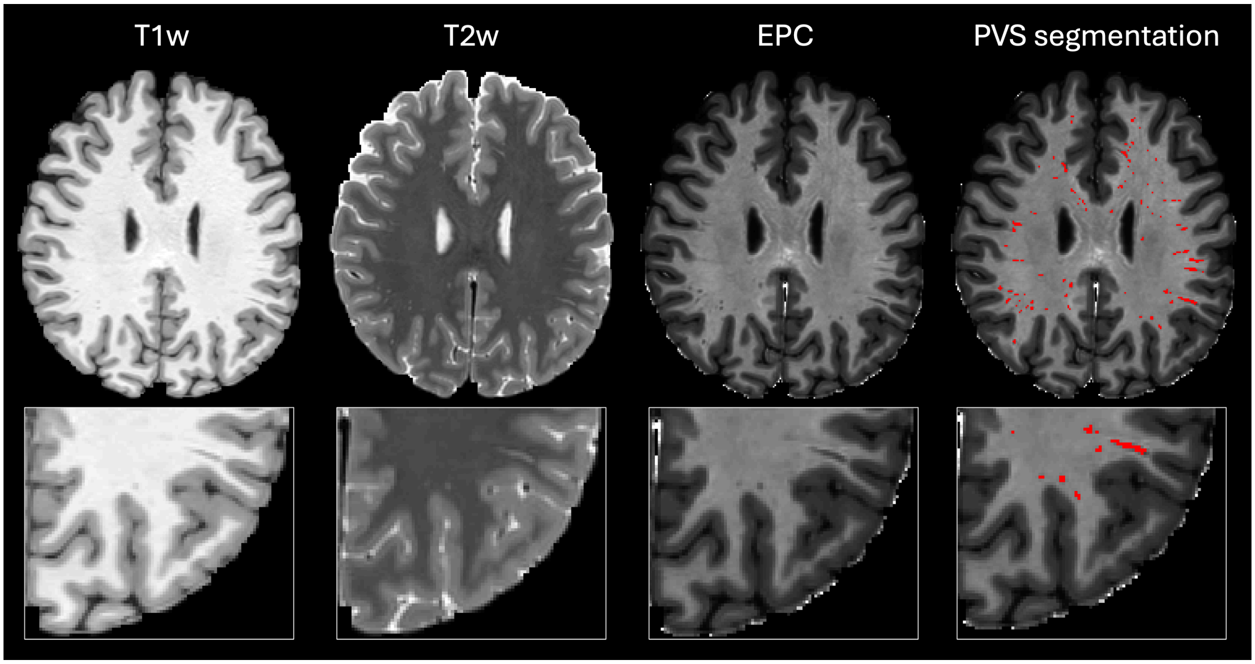


**Supplementary Figure 1. Comparing Enhanced PVS Contrast (EPC) with T1-weighted (T1-w) and T2-weighted (T2-w) images with perivascular spaces (PVS) segmentation.**

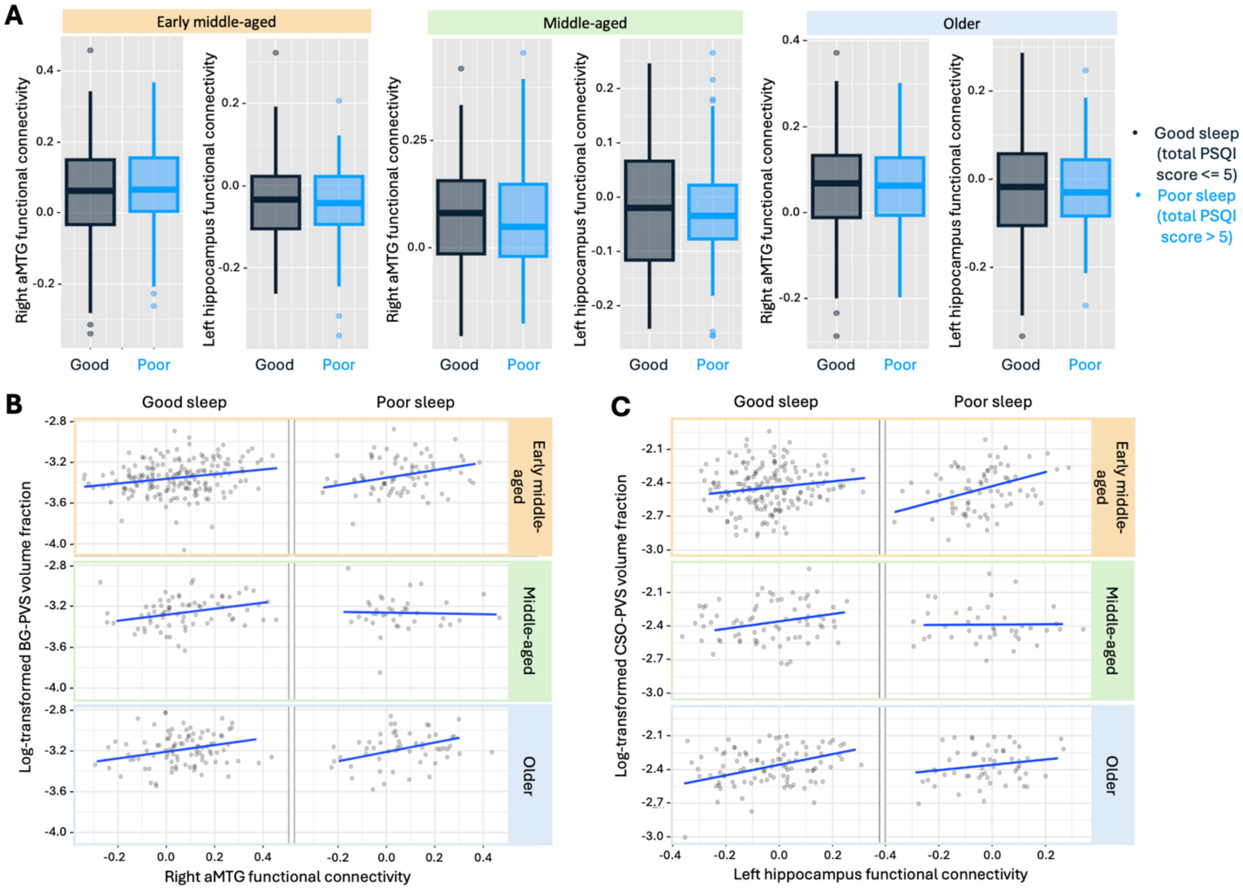


**Supplementary Figure 2. Differences in brain FC between sleep quality groups. (A)** Compared with the good sleep group, the poor sleep group showed no difference of functional connectivity in the right aMTG and right temporal cluster and left hippocampus and frontal cluster (all $p > 0.05$). **(B)** The relationship between BG-PVS and functional connectivity across the different age groups and sleep quality groups. **(C)** The relationship between CSO-PVS and functional connectivity across the different age groups and sleep quality groups.
